# Shark conservation risks associated with the use of shark liver oil in SARS-CoV-2 vaccine development

**DOI:** 10.1101/2020.10.14.338053

**Authors:** Catherine Macdonald, Joshua Soll

**Affiliations:** Field School, 3109 Grand Ave #154, Miami, FL 33133; Rosenstiel School of Marine and Atmospheric Science, University of Miami, 4600 Rickenbacker Causeway, Miami, FL, USA 33149; University of Colorado Boulder, Department of Ecology and Evolutionary Biology

**Keywords:** Squalene, COVID-19, SARS-CoV-2, Shark, Liver oil, Vaccines

## Abstract

The COVID-19 pandemic may create new demand for wildlife-generated products for human health, including a shark-derived ingredient used in some vaccines. Adjuvants are a vaccine component that increases efficacy, and some adjuvants contain squalene, a natural compound derived from shark liver oil which is found most abundantly in deep-sea sharks. In recent decades, there has been growing conservation concern associated with the sustainability of many shark fisheries. The need for a potentially massive number of adjuvant-containing SARS-CoV-2 vaccines may increase global demand for shark-derived squalene, with possible consequences for shark conservation, especially of vulnerable and understudied deep-sea species. A shift to non-animal-derived sources of squalene, which are similar in cost and identical in effectiveness, or an emphasis on increasing traceability and sustainability of shark-derived squalene from existing well-managed fisheries, could better support conservation and public health goals.

## 1. Introduction

Shark populations have historically been exploited for meat, fins, leather, and products derived from their liver oil, including vitamin A and additives to topical cosmetic formulas and lotions (Dent and Clarke, 2015; Lozano-Grande et al., 2018). Some historic shark liver oil fisheries have been associated with intense, short-lived periods of exploitation contributing to stock collapses (Stevens et al., 2000; Kyne and Simpfendorfer, 2007; Anderson and Waheed, 1999; Ali and Sinan, 2014). Sharks, particularly deep-sea species, tend to grow slowly and produce relatively few young, leaving them especially susceptible to overexploitation (Simpfendorfer and Kyne, 2009). Squalene is a natural polyunsaturated hydrocarbon compound (C_30_H_50_) found in many living organisms that can be refined from shark liver oil. This compound (and its more stable, hydrogenated derivate, squalane) have shown genuine potential to contribute to human health, as an ingredient in vaccines and other pharmaceuticals and dietary supplements (Kim and Karadeniz, 2012).

Many shark species, particularly deep-sea sharks, have relatively large livers (up to 20% of animal weight) which serve as a buoyancy control system and critical energy reserve (Vannuccini 1999; Abel and Grubbs 2020). A significant portion of liver weight is made up of liver oil (10-70%; Nichols et al., 2001), and between 15 and 82% of liver oil is squalene (Deprez et al., 1990; Bakes & Nichols 1995). Although squalene is produced by a range of animals and plants, shark liver oil has historically been a preferred commercial source based on availability and high yields relative to most plant-derived sources.

Squalene can be extracted directly from shark liver oil at high purity (>98%) in approximately 10 hours, using a single distillation process in a vacuum at 200-230° C, a faster process producing greater yields than plant-based alternatives (Camin et al., 2010). Sealed and protected from oxygen and light, squalene has an approximately two-year shelf life, making stable rates of ongoing production important to availability (Camin et al., 2010). Plant-derived squalene can also be refined to a high level of purity for medical applications, and is made up of C_30_H_50_ molecules chemically identical to those from shark-derived sources. Plant-derived squalene has been shown to perform comparably to shark-derived squalene as an ingredient in vaccine adjuvants (Brito et al. 2011). Accordingly, the FDA is agnostic on the source of squalene and the *Current Good Manufacturing Practice for Finished Pharmaceuticals* only requires that lots of C_30_H_50_ be “tested for conformity with all appropriate written specifications for purity, strength, and quality” (2019). Chemical vendors listed on the NIH National Library of Medicine for squalene do not differentiate by squalene origin but instead identify the compound by Chemical Abstract Services (CAS) number 7683-64-9, meaning there are no legal or administrative barriers preventing a shift to plant-derived sources of squalene (National Center for Biotechnology Information, 2020).

Shark-derived squalene is a popular ingredient in topical cosmetic formulas and lotions, generating a significant portion of global demand for shark liver oil (Lozano-Grande et al., 2018; Ducos et al., 2015). In recent years, however, consumers and non-profit organizations in the United States and Europe have exerted pressure on the cosmetics industry to transition to plant-derived squalene (Consumers unaware of… 2013; Shark Free). Independent testing by the French non-profit organization Bloom determined most beauty products (>90% of those tested) sold in Europe or the United States no longer contain shark-derived ingredients, although shark-derived squalene remains in common use in beauty products elsewhere (Ducos et al., 2015). More recent studies have demonstrated the presence of shark DNA, including from Threatened and Endangered species, in some US-sold beauty products as well (Cardeñosa et al., 2017; Cardeñosa, 2019).

### 1.1 Squalene in vaccines

Squalene is used as an adjuvant, a component of a vaccine that enhances efficacy by increasing human immune response and the potency of antigens, while promoting antigen transport into lymph nodes and uptake in the immune cells (Brito and O’Hagan, 2014). The inclusion of adjuvants also reduces the volume of antigen needed per vaccine dose through the process of “antigen sparing,” maximizing rates of vaccine production (Tang et al.. 2015; Schmidt et al. 2016). Although the primary global demand for shark liver oil in recent years appears to be for use in cosmetics and lotions (Vannuccini 1999; Lozano-Grande et al., 2018), the role of shark-derived squalene in vaccine production may represent a potential threat to some vulnerable shark populations, given widespread efforts to rapidly develop and disseminate vaccines in response to the COVID-19 pandemic and the patchwork management approach to global shark fishing limits (Ahn et al., 2020; Hodgson, 2020; Davidson et al. 2016).

As an ingredient in adjuvants, squalene has been found to be safe and effective in vaccines for the treatment of viruses including H7N9 and H7N7 (Wu et al., 2014), H1N1 (Vesikari et al., 2012), MERS-CoV (Zhang et al. 2016), and SARS-CoV (Stadler et al., 2005). MF59, an adjuvant containing shark-derived squalene, has been used commercially in influenza vaccines (Schultze et al. 2008; Panatto et al. 2020) and used or tested in previous vaccines for other coronaviruses (Stadler et al., 2005; Zhang et al., 2016).

A few of the potential SARS-CoV-2 vaccines currently in testing include adjuvants MF59 or AS03, which are known to contain shark-derived squalene (COVID-19 Update 2020; GlaxoSmithKline aims… 2020, WHO, 2020). Per dose, MF59 contains 9.75 mg and AS03 contains 10.68 mg of squalene (Jackson et al., 2015). Some potential vaccines are being manufactured in bulk before the completion of large-scale safety and efficacy trials (U.S. to Stockpile… 2020), suggesting that the total number of manufactured doses of various potential vaccines may be far higher than the number ultimately administered. It also remains unknown how frequently revaccination would be necessary if a safe and effective vaccine becomes available (COVID-19 Could Become… 2020; Ellis, 2020; Vaccine companies… 2020), so the scale of possible future demand for squalene as an ingredient in vaccine adjuvants is large but difficult to fully or accurately assess.

Deep-sea sharks are valued in the shark liver oil trade because they offer greater volumes of liver oil than other species. Relatively little is known about population structure, reproduction, and habitat use of many of these species due to the logistical difficulties of conducting research in deep-sea habitats and a generally low research and management priority (e.g., Neiva et al., 2006; Kyne and Simpfendorfer, 2007; Veríssimo et al., 2011). Most elasmobranchs are slow to mature and reproduce, and deep-sea sharks are particularly so, showing population increase rates of less than half those of continental shelf and pelagic shark species relying on shallower habitats (Simpfendorfer and Kyne, 2009). Squaloid sharks (species found in order squaliformes) are a preferred source of liver oil and squalene, and include species which are among the slowest growing and latest maturing sharks known (Musick, 2005). Shark reproductive rates and recovery potential generally decline with increasing depth, and overfished populations of deep-sea shark species may require centuries to recover (Simpfendorfer and Kyne, 2009). Evidence suggests that even low rates of deep-sea shark exploitation (incidental or targeted) is likely to quickly deplete populations (Simpfendorfer and Kyne, 2009). For these reasons, deep-sea sharks have been identified as a conservation priority grouping (Dulvy et al., 2014).

Assessment of the global conservation status of sharks is complicated by poor catch reporting, challenges with species-level identification, and a lack of baseline data. The reported annual catch of sharks likely substantially underestimates actual total catch (Worm et al., 2013; Dulvy et al., 2014). Total catch has been estimated at approximately 100 million animals annually, with an estimate range of 63 to 273 million (Worm et al., 2013). It is particularly difficult to assess rates of capture for species targeted for products like shark liver oil, for which there is relatively little detailed production or trade information available (Fowler et al., 1997; Vannuccini, 1999).

The global trade in shark liver oil is small compared to that in shark fins or meat, and despite the vulnerability of some target species to overfishing has received relatively little media (Shiffman et al., 2020) or scientific attention (Dent and Clarke, 2015). Although there are fisheries in which deep-sea sharks are targeted (see, e.g., Simpfendorfer and Kyne, 2007), liver oil may also be generated opportunistically from incidental catches in fisheries predominantly targeting other fish, or targeting sharks for other products (e.g., meat, fins). In at least some cases, however, the export value of shark liver oil exceeds that of shark meat, and in some locales the available supply of shark liver oil is insufficient to meet processors’ demand for raw materials (Dent and Clarke, 2015). A lack of traceability presents a further challenge in assessing the trade in shark liver oil, as the United Nations Food and Agriculture Organization (FAO)’s *Codex Alimentarius* allows for products like squalene to be labelled as originating in the countries in which they were processed even when ingredients (e.g., the liver oil from which the squalene was derived) were produced elsewhere (FAO, 2001).

This study provides a basis for beginning to evaluate the range of potential conservation effects of increasing demand for shark-derived squalene as an ingredient in vaccine adjuvants.

## 2. Materials and methods

Based on a review of available scientific and management literature, 133 species were identified which are known to be involved in the liver oil trade (Appendix 1), including many that are partially reliant on deep-sea habitat >200 m (n=83) or are found exclusively in the deep-sea (n=21). Additional data on the conservation status and population trends of elasmobranch species, primarily sharks, identified as being involved in the liver oil trade were compiled from their most recent IUCN Red List assessments. Available current trade data on the shark liver oil trade was downloaded from the UN FAO FishStat Database (FAO, 2020). Commercial sources of wholesale quantities of squalene were identified through internet searches and through direct contact with wholesalers to collect pricing, origin, and availability information for both plant- and shark-derived squalene and squalane (its more stable, hydrogenated derivate).

## 3. Results

Across elasmobranch species identified as being exploited for shark liver oil, 50% had not been assessed by the IUCN in at least ten years, and 30% were considered Data Deficient (Figure 1; an IUCN Shark Specialist Group reassessment of elasmobranch species is currently underway and expected to be completed in 2020 (Dulvy, 2018)). One-third of species identified as part of the shark liver oil trade are classified as threatened (Vulnerable, Endangered, or Critically Endangered) according to IUCN Red List criteria (Figure 2). Population trends for 56% of these species are unknown, and 34% are assessed as showing a decreasing trend (Figure 3).

**Figure 1.**
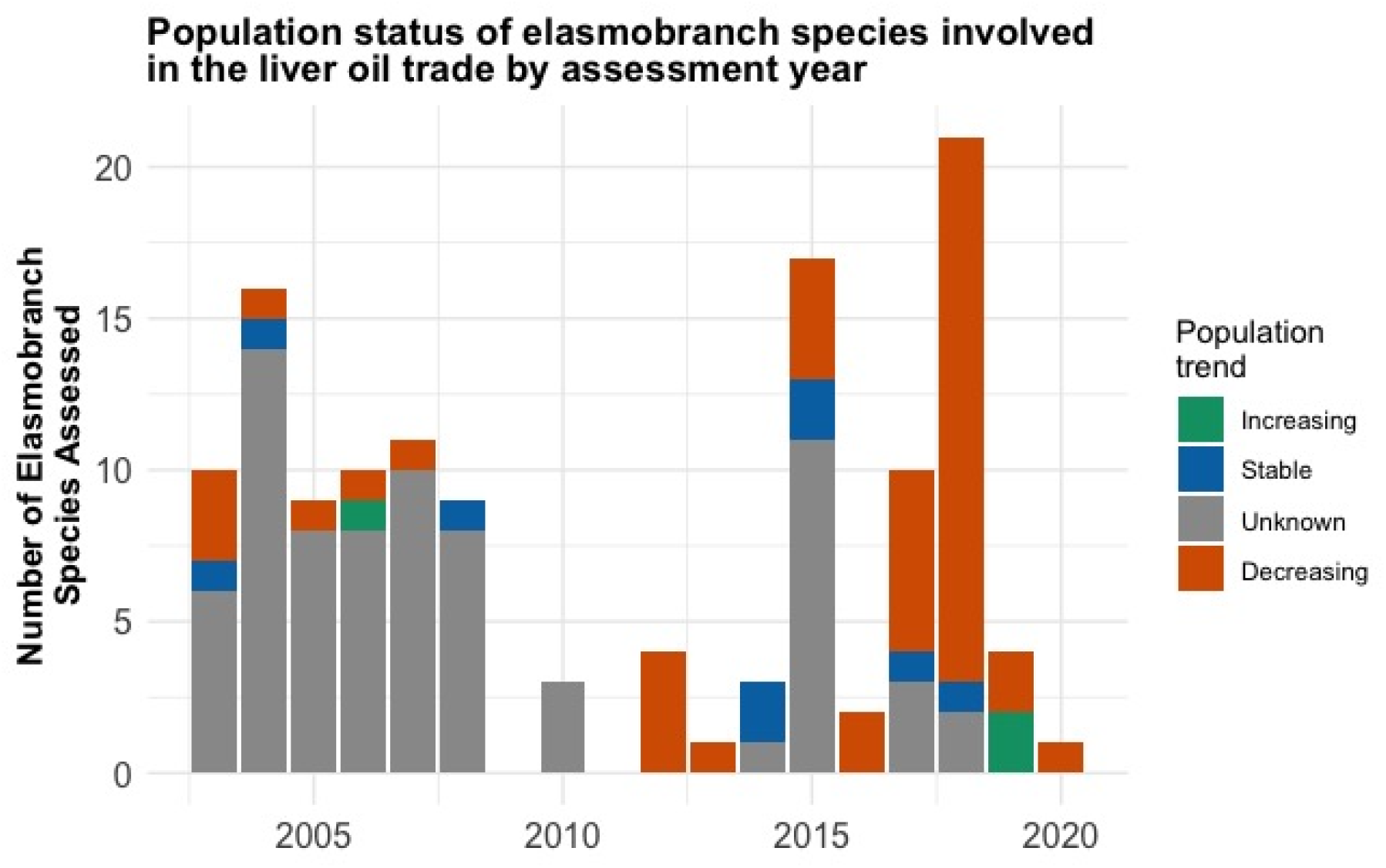
Most recent IUCN Red List assessments for assessed elasmobranch species (n=131) reported in the liver oil trade. The earliest assessment of a species identified as traded for liver oil took place in 2003.

**Figure 2.**
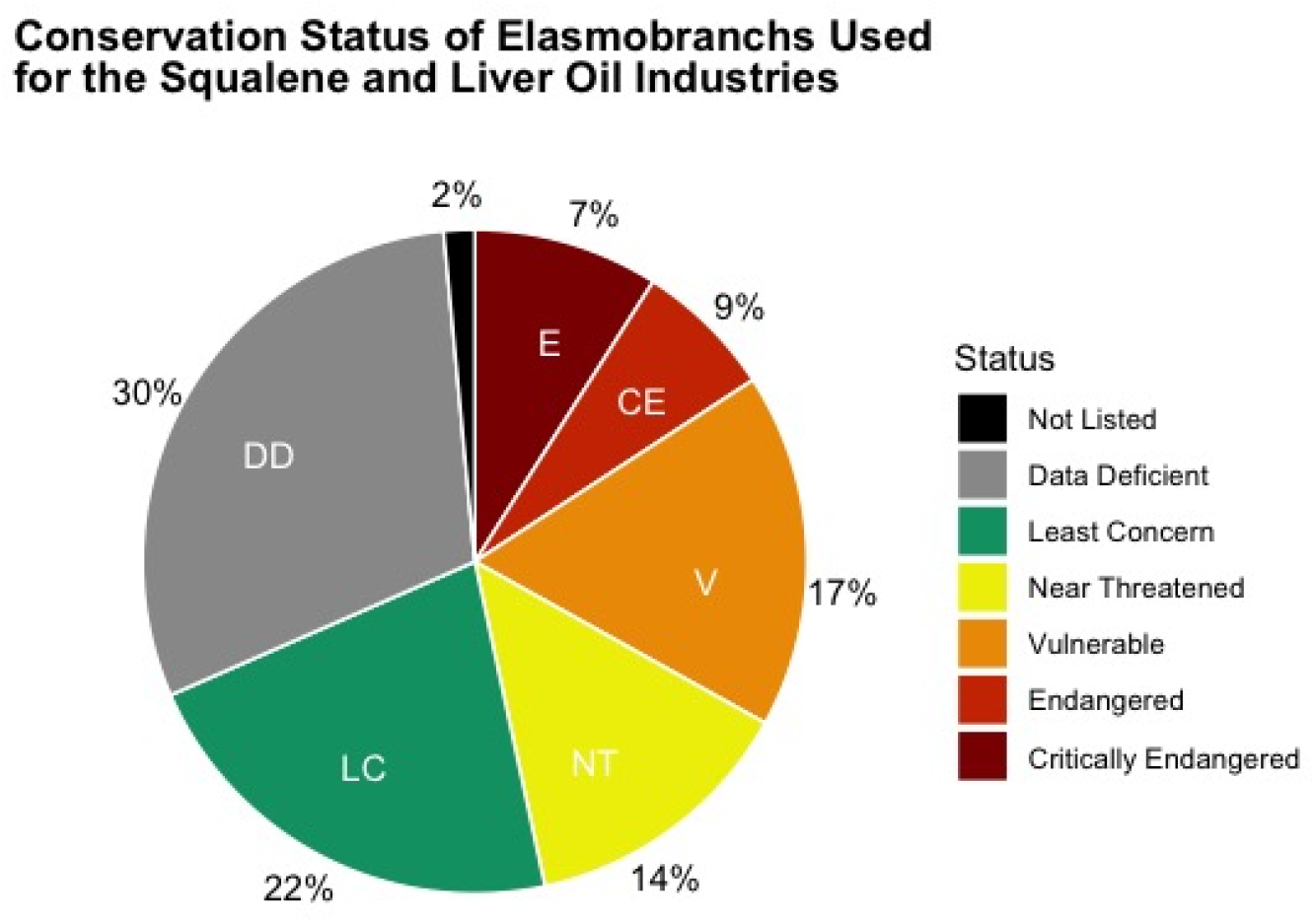
IUCN Red List conservation status of elasmobranch species reported in the liver oil trade.

**Figure 3.**
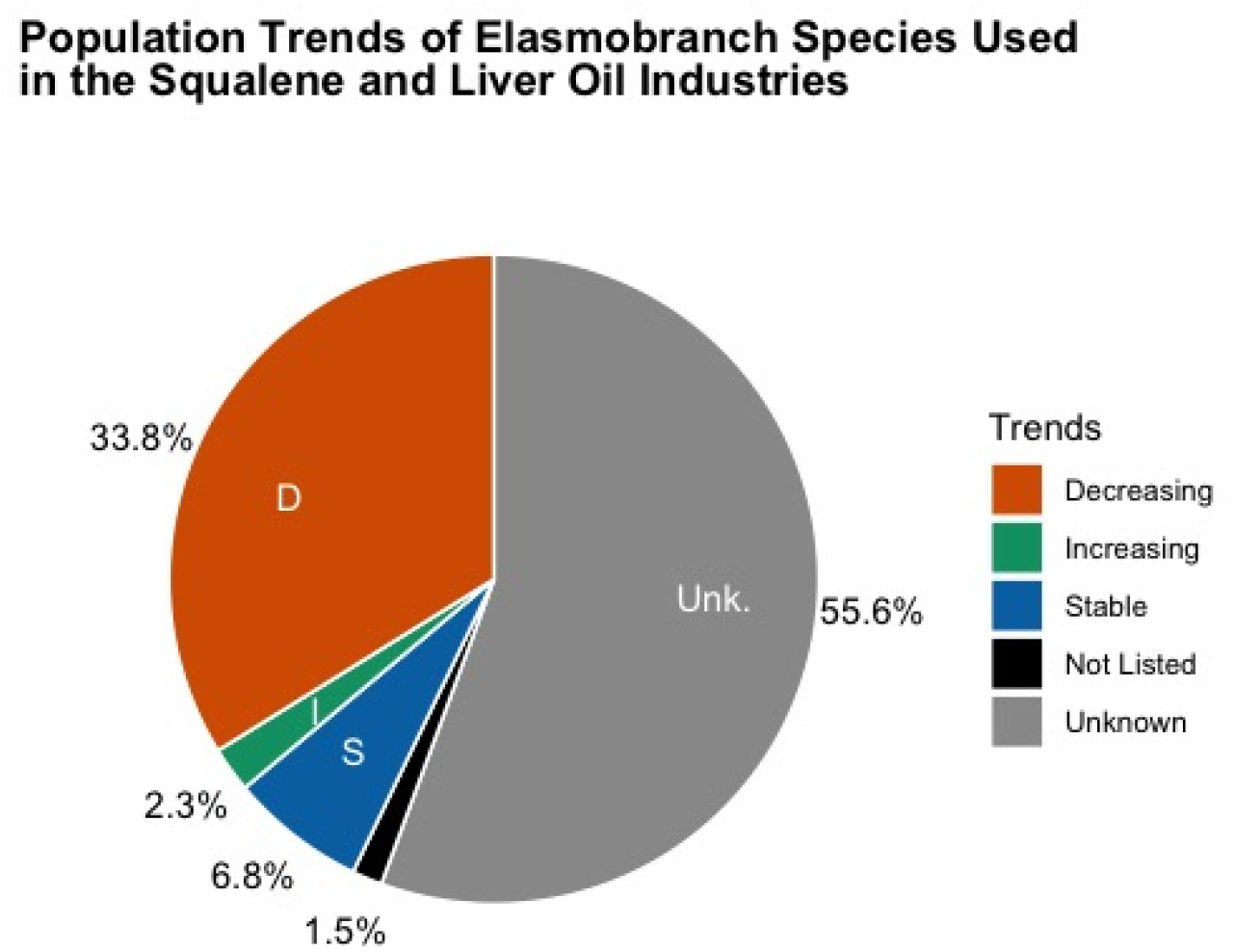
IUCN Red List population trends of all elasmobranch species reported in the liver oil trade.

The most recent FAO data available (2018) showed an increase in reported import and processed production of shark liver oil, with trade volumes reaching 752 tons, the largest reported volume in decades (Figure 4; from 2000-2017, average reported trade volume was under 200 tons annually; Hareide et al., 2007). Despite this increase, the total value of the shark liver oil trade was reported at 553 000 USD in 2018, the lowest value reported since 1987 (mean annual value from 2000-2017, 1 954 000 USD; Hareide et al. 2007; FAO, 2020). Moderate assumptions about oil yield suggest the range of individual animals in the reported global liver oil trade in 2018 falls between 694 848 (large sharks) and 16.35 million (small sharks), with 1.8 million (based on a 20.8 kg median unidentified shark weight; Worm et al. 2013) as the best supported rough estimate (for yield assumptions see Table 1).

**Figure 4.**
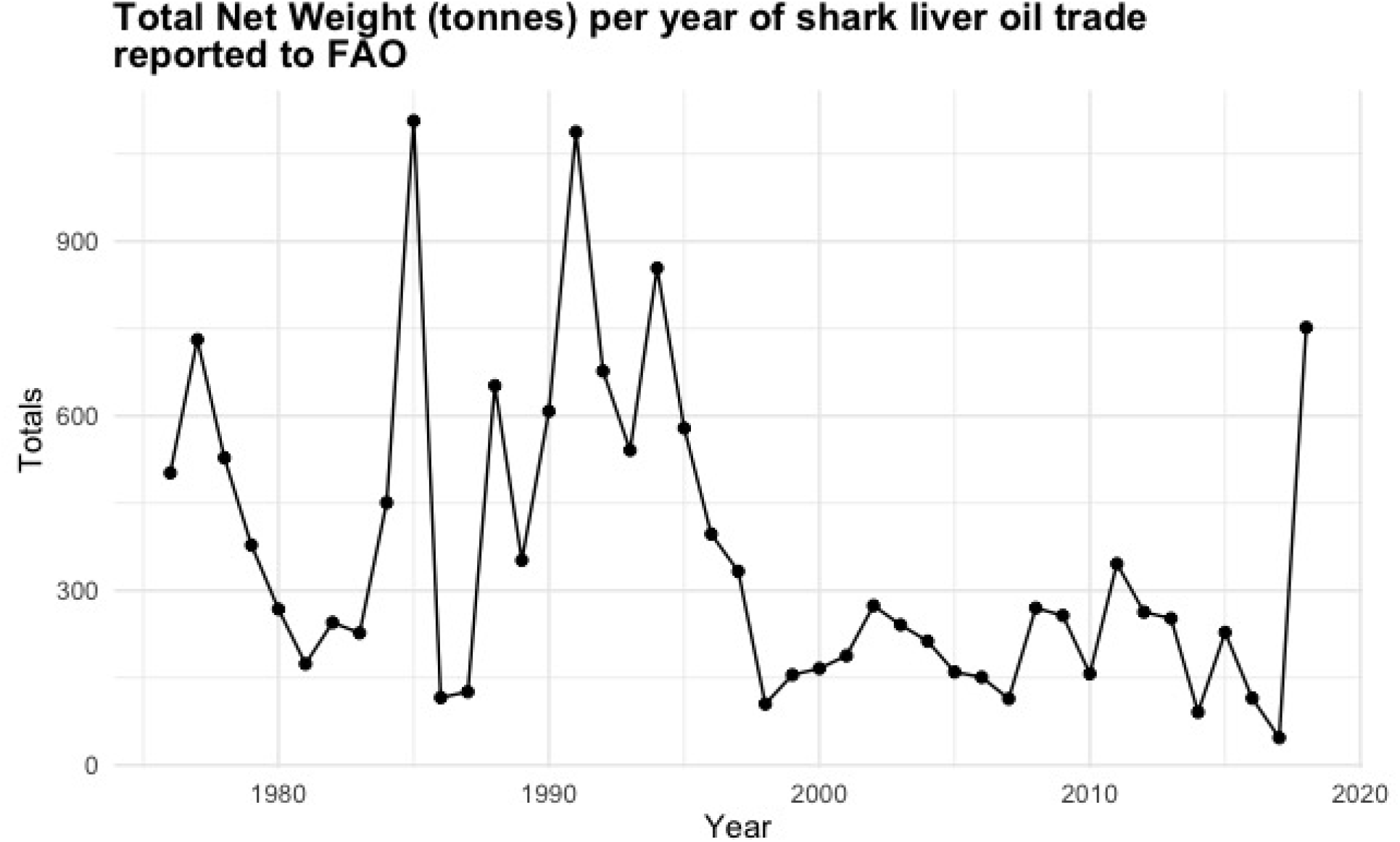
The total reported net weight (tonnes) of annual trade in shark liver oil reported to FAO. In 2018, two countries reported trade: imports of 33 tonnes (Republic of Korea) and processed production of 719 tonnes (Senegal).

**Table 1.** Estimated number of sharks needed to produce two vaccine doses for the global human population under a range of yield assumptions.

| Shark weight (kg) | Liver weight | Liver oil weight | Squalene weight | Total sharks* |
| --- | --- | --- | --- | --- |
| Best case |  |  |  |  |
|  | <u>20%</u> | <u>70%</u> | <u>82%</u> |  |
| 2.3 kg (small shark) | 0.46 | 0.32 | 0.26 | 576,049 |
| 5.6 kg (deep-sea shark) | 1.12 | 0.78 | 0.64 | 236,592 |
| 20.8 kg shark (median) | 4.16 | 2.91 | 2.39 | 63, 693 |
| 46.2 kg (large shark) | 9.24 | 6.47 | 5.30 | 28,678 |
| Moderate case |  |  |  |  |
|  | <u>10%</u> | <u>40%</u> | <u>50%</u> |  |
| 2.3 kg (small shark) | 0.23 | 0.09 | 0.05 | 3,306,522 |
| 20.8 kg shark (median) | 2.08 | 0.83 | 0.42 | 365,385 |
| 46.2 kg (large shark) | 4.62 | 1.85 | 0.92 | 164,610 |
| Worst case |  |  |  |  |
|  | <u>3%</u> | <u>10%</u> | <u>15%</u> |  |
| 2.3 kg (small shark) | 0.069 | 0.007 | 0.00104 | 146,956,522 |
| 20.8 kg shark (median) | 0.624 | 0.062 | 0.00936 | 16,239,316 |
| 46.2 kg (large shark) | 1.386 | 0.139 | 0.02079 | 7,316,017 |

\* Based on a maximum demand estimate, assuming use of MF59 adjuvant containing 9.75 mg of squalene per dose to provide 2 vaccines to 7.8 billion people, total squalene demand for vaccines (drawn from existing or new fisheries) would be 152,100 kg.

Data collected on the price of shark- and plant-derived squalene showed no clear differences, though products listed as both shark- and plant-derived were commonly available in the wholesale market (Table 2). There was significant variability in price across individual sellers (ranging from $20-260/kg for squalene and $10-99.21/kg for squalane). The mean and median price for plant-derived squalene was $40, and the mean price for shark-derived squalene was $45.75 with a median of $48.5/kg (excluding the outlier of $260; including it, the mean price for shark-derived squalene was $76.36, with a median price of $54). The mean price for plant-derived squalane was $53.16/kg (median $48.65/kg), while shark-derived squalane was $45/kg (median $45/kg). These products were identified as being sold from China (67% of products) and the United States (33% of products). Based on these numbers, the estimated potential commercial value of squalene from a single shark ranges from $0.05 (small shark/low yield) to $242.65 (large shark/high yield).

**Table 2.** Current listed wholesale prices of shark- and plant-derived squalene and squalane by kg in USD.

| Squalene |  |  |  |
| --- | --- | --- | --- |
| <u>Source</u> | <u>Price/Kg</u><br><u>(USD)</u> | <u>Sold From</u> | <u>Manufacturer</u> |
| Shark | \$39-47 | China | Xi'an Rozen Biotechnology Co., Ltd |
| Shark | \$25 | China | Xa Bc-Biotech Co., Ltd |
| Shark | \$54 | China | Wuhan Hengheda Pharm Co., Ltd |
| Shark | \$260 | China | Haihang Industry Co., Ltd |
| Shark | \$35-80 | China | Xi'an Sheerherb Biological<br>Technology |
| Shark | \$20-60 | China | Shandong Zesheng Chemical Co., Ltd |
| Shark | \$55 | China | Shaanxi Phoenix Tree Biotech Co.,<br>Ltd |
| Olive Oil | \$20-60 | China | Suzhou Manson Biotech Co., Ltd |
| Soybeans | \$20-60 | China | Suzhou Manson Biotech Co., Ltd |
| Squalane |  |  |  |
| <u>Source</u> | <u>Price/Kg</u><br><u>(USD)</u> | <u>Sold From</u> | <u>Manufacturer</u> |
| Shark | ~\$40 | USA | Jedwards International INC. |
| Shark | \$50 | China | Xa Bc-Biotech Co, Ltd |
| Olive Oil | \$44.89 | USA | Cocojojo Beauty |
| Olive Oil | \$63.99 | USA | Organic Gold |
| Olive Oil | \$10-20 | China | Shandong Zhonglan Chemical Co.,<br>Ltd |
| Olive Oil | \$40-60 | China | Xi'an Geekee Biotech Co., Ltd |
| Olive Oil | ~\$51.75 | USA | Jedwards International INC. |
| Olive Oil | ~\$47.30 | USA (Origin Italy,<br>Spain) | Blossom Bulk Ingredients |
| Neossance (Sugarcane) | \$99.21 | USA | HBNO |

Given that each vaccine dose requires approximately 10mg of squalene, the mean squalene cost-per-vaccine-dose for shark-derived squalene would be $0.0004575 USD, and for plant-derived, $0.0004 USD—a price difference of $0.0000575 per dose.

## 4. Discussion

These results highlight the extent to which liver oil fisheries could affect shark species of conservation concern, and the potential difficulty of detecting these effects because of a lack of information on the volume of liver-oil-associated catch and the absence of population status information for many deep-sea species. The life history characteristics of deep-sea sharks (Simpfendorfer and Kyne, 2009), insufficient restrictions on exploitation, and declines in availability of shark-derived squalene over time (Sibuyo et al., 2017) suggest that the trade in shark liver oil, while currently small, has potential to disproportinately harm specific vulnerable elasmobranch species and populations.

Estimating potential conservation effects is further complicated by the fact that shark species vary greatly in the percentage of their body weight comprised of the liver (from at least 2.9-20%; Vannucini, 1999; Abel and Grubbs, 2020); in the amount of liver weight made up of oil (ranging from at least 10-70%; Nichols et al., 2001); and in the yield of squalene from extracted liver oil (a range of at least 15-82%; Deprez et al., 1990; Bakes & Nichols 1995). Evidence further suggests animal size, sex, and seasonal and regional factors can also affect yields (Kreuzer and Ahmed, 1978; Nichols et al., 2001). Accordingly, it is impossible to calculate an exact effect of increased vaccine-related demand for shark-derived squalene on shark populations.

We know that fisheries primarily targeting sharks for liver oil exist (e.g., Kyne and Simpfendorfer, 2007), and that, in many cases, multiple products may be marketed from a single shark (e.g., meat, fins, liver oil, cartilage). Data are not available to assess the volume of liver oil generated globally by directed versus incidental fishing. The effect of increased demand for shark-derived squalene is highly dependent on whether it increases targeting of oil-rich deep-sea sharks, or simply drives increased processing and use of the livers of sharks currently taken for other purposes. Thus, conservation effects of vaccine-related demand for squalene could range from minimal (assuming demand is met by more efficient use of individuals already being landed in well-managed fisheries) to catastrophic for individual species (if demand drives the creation of new targeted fisheries for vulnerable deep-sea species in the absence of limits on fishing and trade).

The future availability of shark squalene will likely be constrained by shark population declines and by regulatory efforts to conserve sharks, suggesting the prudence of shifting to more stable and sustainable sources. A vaccine supply chain dependent on shark fisheries is subject to disruption if targeted shark populations collapse or new protections are introduced. Plant-derived alternatives to shark-derived squalene are more environmentally sustainable and readily available. Although in the past shark-derived squalene was reported to be significantly less expensive than plant-derived (Camin et al., 2010), current costs are similar. Non-animal-derived squalene may even be safer for use in some health-related applications because of the reduced risk of contamination with persistent organic pollutants, including polychlorinated biphenyls and organochlorine insecticides, high levels of which have been found in nutritional supplements made from shark liver oil (Rawn et al., 2009).

Despite generally comparable costs between shark- and plant-derived squalene, and potential advantages in long-term availability and contaminant levels, non-animal-derived squalene does not appear to be in current use in vaccine production. Consumer and activist pressure could be effective in encouraging pharmaceutical companies to transition to non-animal-derived squalene in adjuvants, for testing for other medical uses, and in nutraceutical products. Demand-driven increases in production of pharmaceutical grade plant-derived alternatives may also incidentally support reduction or elimination of shark-derived squalene in cosmetic formulations. Increased accountability, including product testing to confirm plant origins and transparency within supply chains, would be vital to a transition away from reliance on shark-derived squalene.

## 5. Conclusion

The mean difference in cost per dose of a potential SARS-CoV-2 vaccine containing squalene derived from plants instead of shark liver oil is −$0.0000575, based on currently available wholesale price information. While commercially available wholesale squalene may not meet the standard of purity needed for pharmaceutical applications, given these values it would cost $20,125.00 less to generate the 350 million doses needed to vaccinate the U.S. population with plant-rather than shark-derived squalene. Doing the same for the global population of 7 billion would save $402,500.00 over use of shark-derived squalene (without accounting for the effects high demand might have on price or availability).

In addition to similarities in cost, plant-derived sources of squalene have been shown to work comparably as an ingredient in adjuvants, and should not require a new approval process for use because they are chemically identical. Therefore, a shift to non-animal-derived squalene is highly feasible without disrupting efforts to rapidly develop a vaccine. A coordinated commitment by the medical sector to transition to non-animal-derived squalene would support both environmental sustainability and public health goals.

## Acknowledgments

We are indebted to Dr. Julia Wester, Dr. David Shiffman, Sonja Fordham, Brandon Theiss, and Dr. Brynn Devine for reading and providing helpful feedback on earlier drafts. Author JS would also like to thank Stefanie Brendl, Alexia Skrbic, and Laurel Irvine for encouraging him to pursue research opportunities associated with this project as part of his internship with Shark Allies.

## Appendix A: Elasmobranchs reported to be used in the liver oil and squalene trades, including IUCN conservation status and population trend assessment information

**Table A.1.**
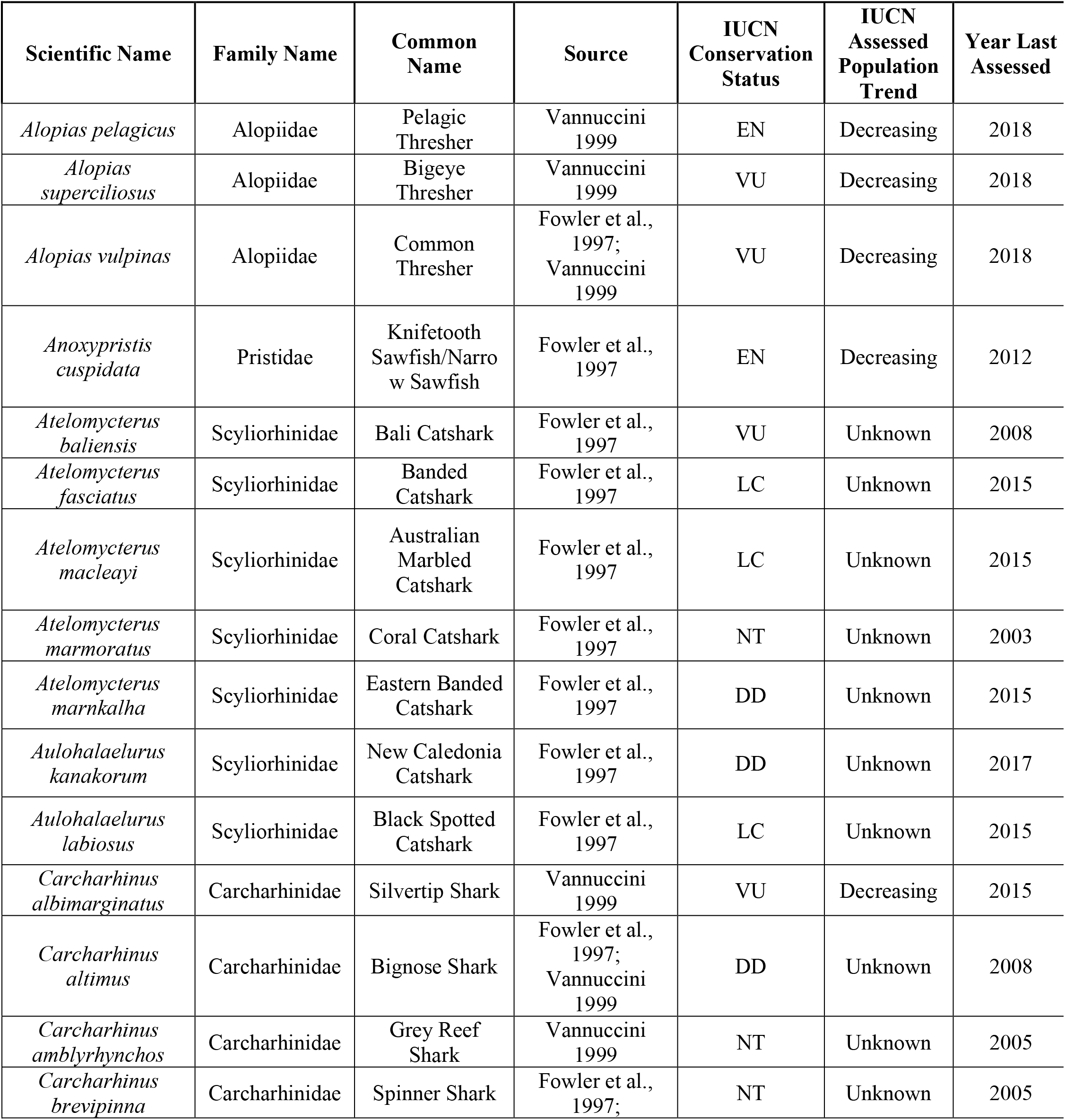
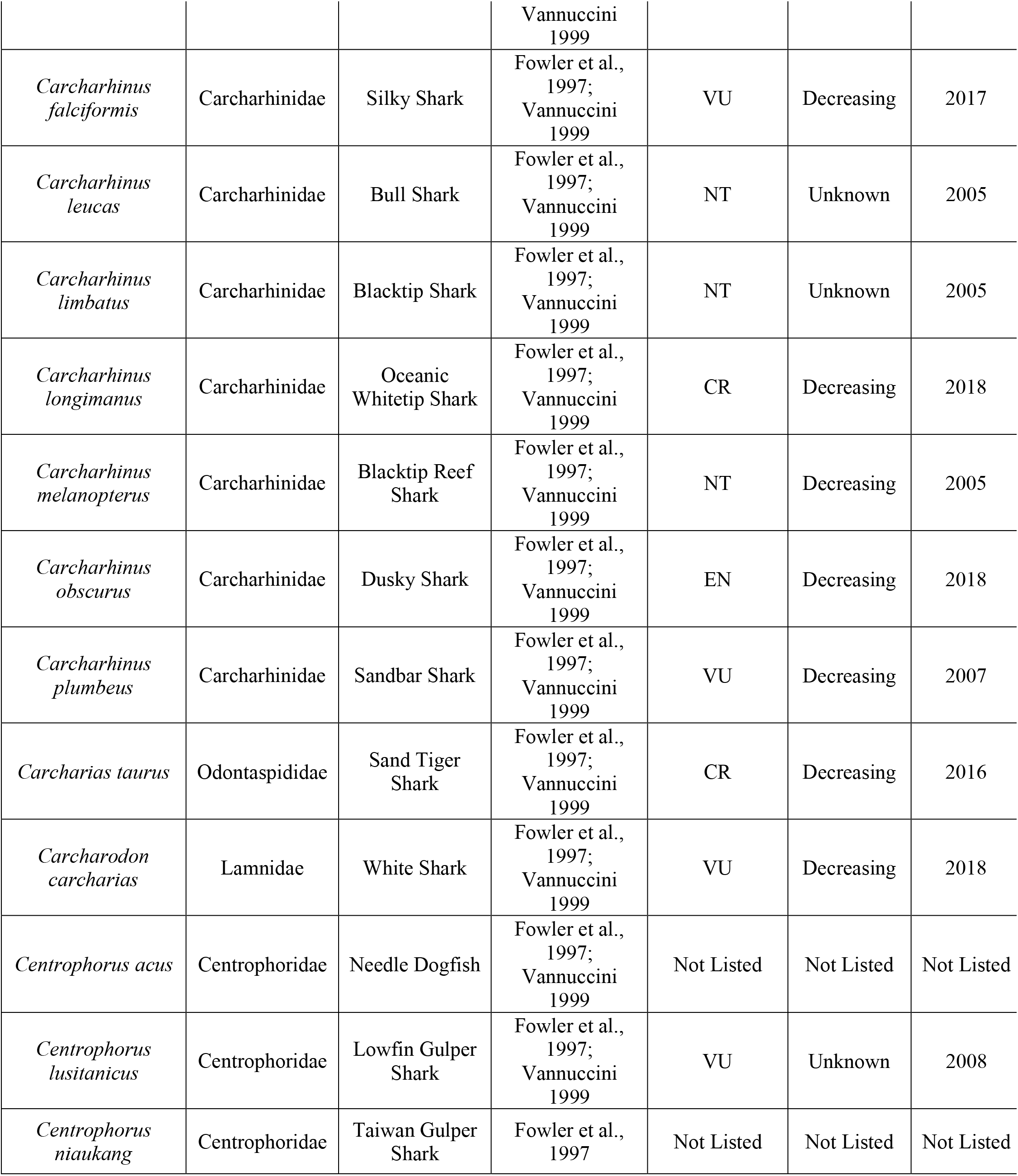
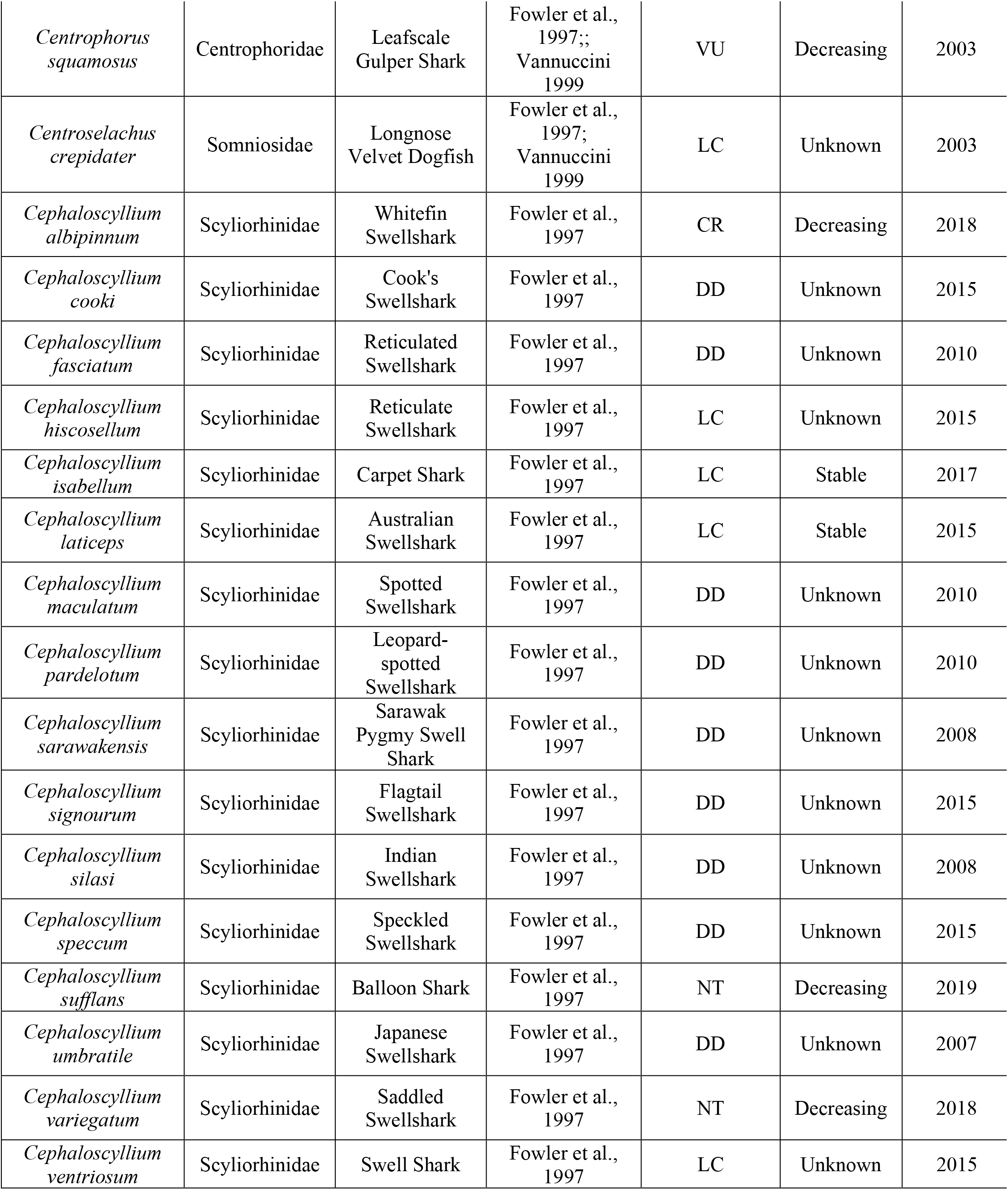
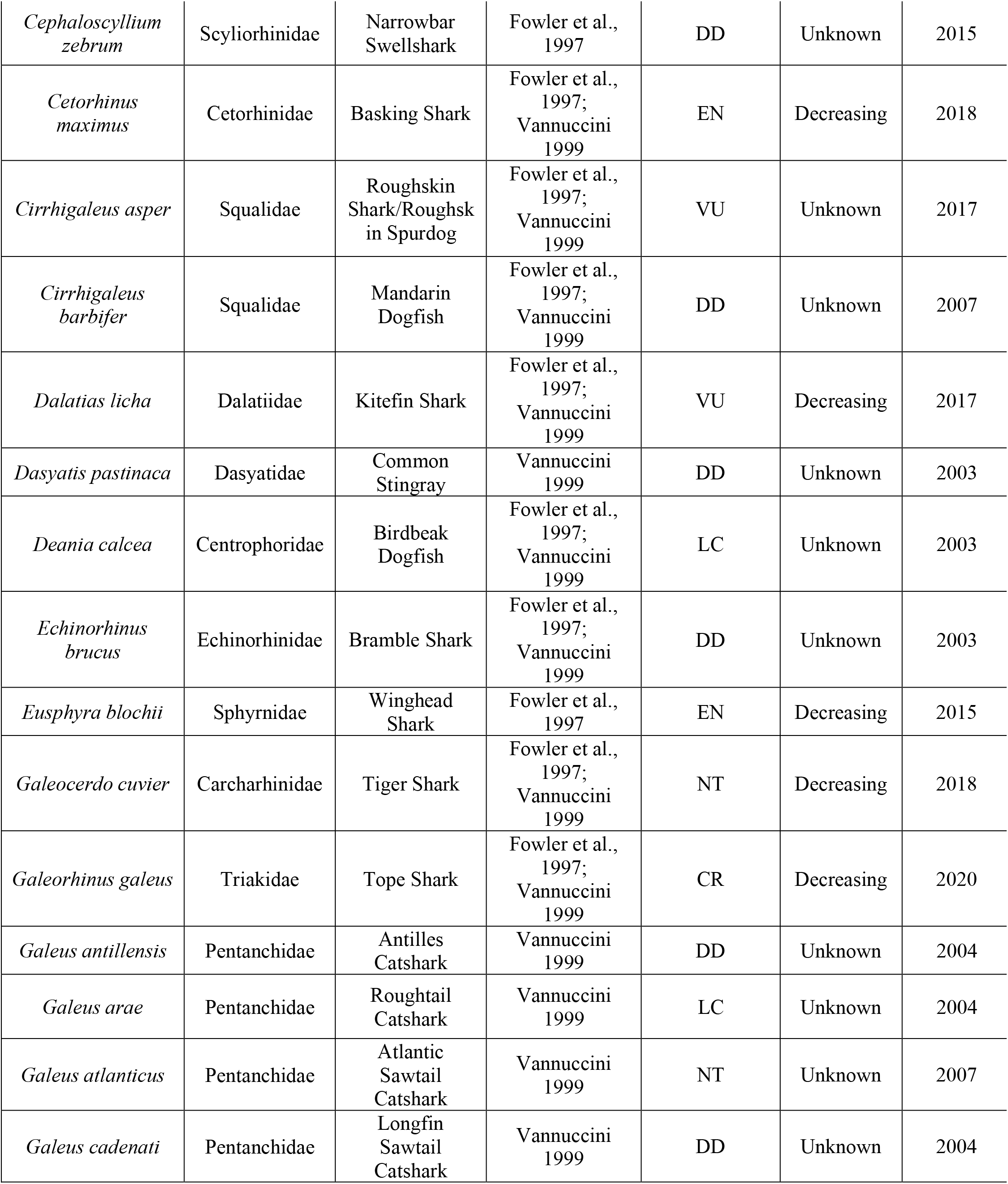
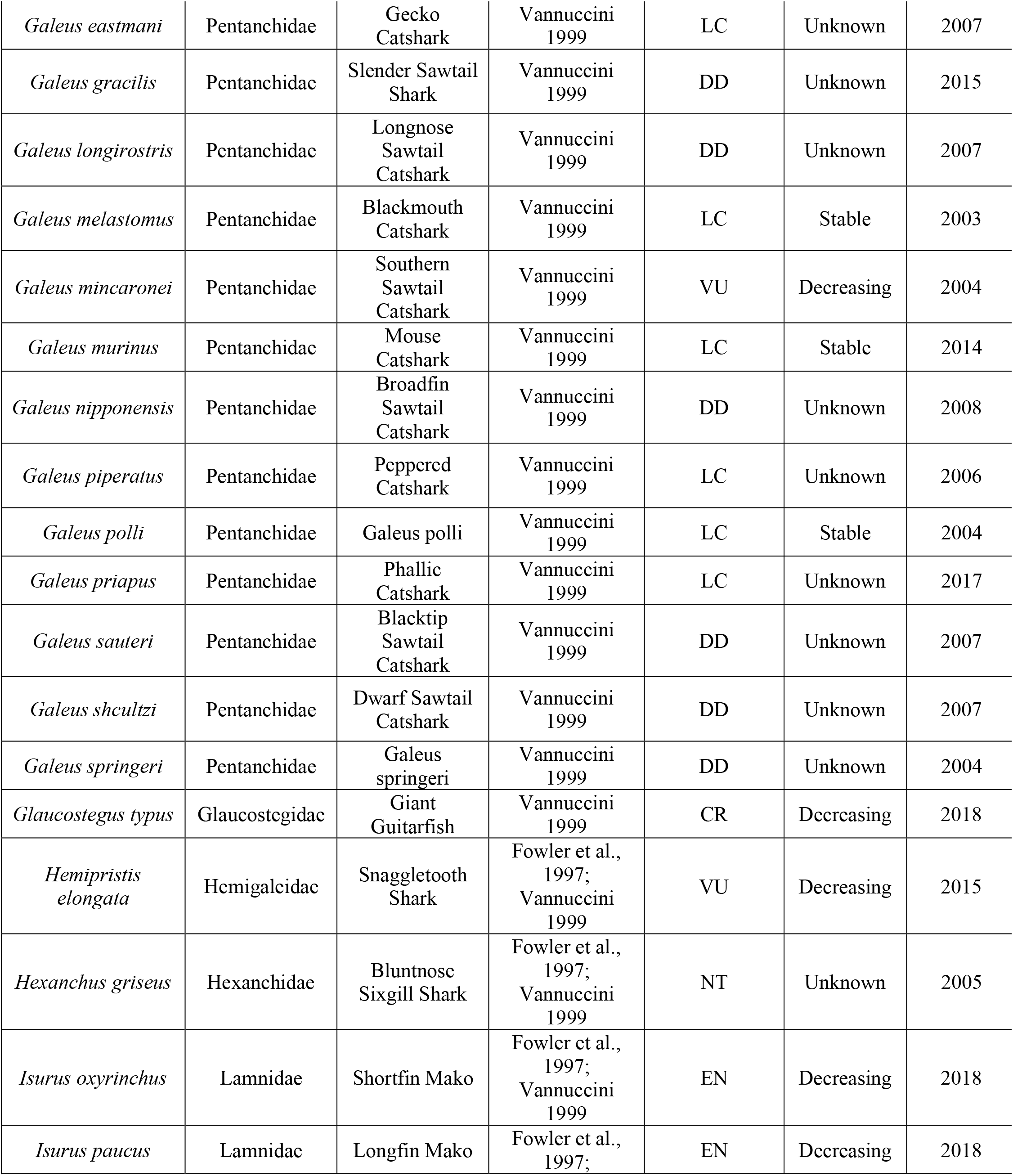
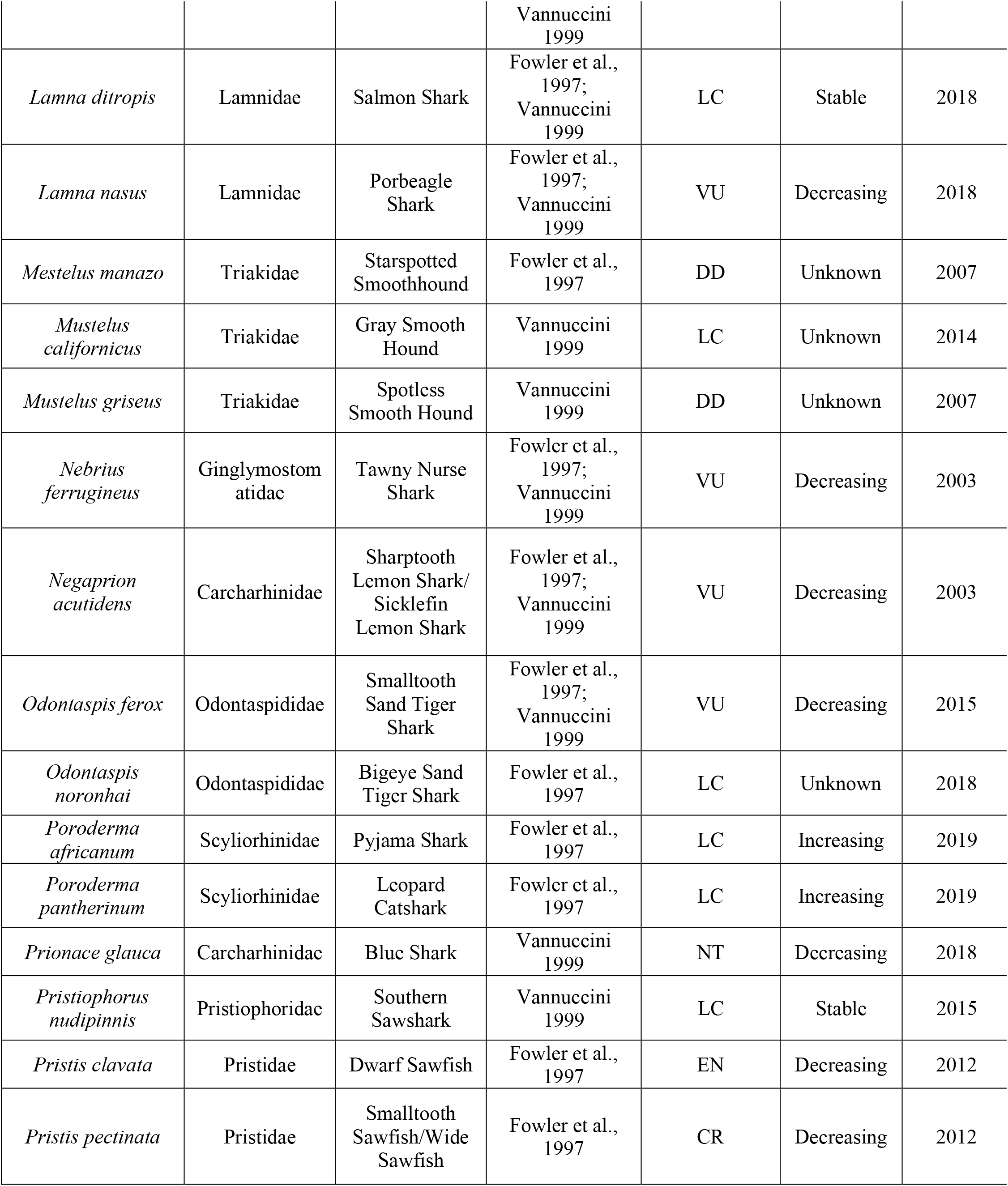
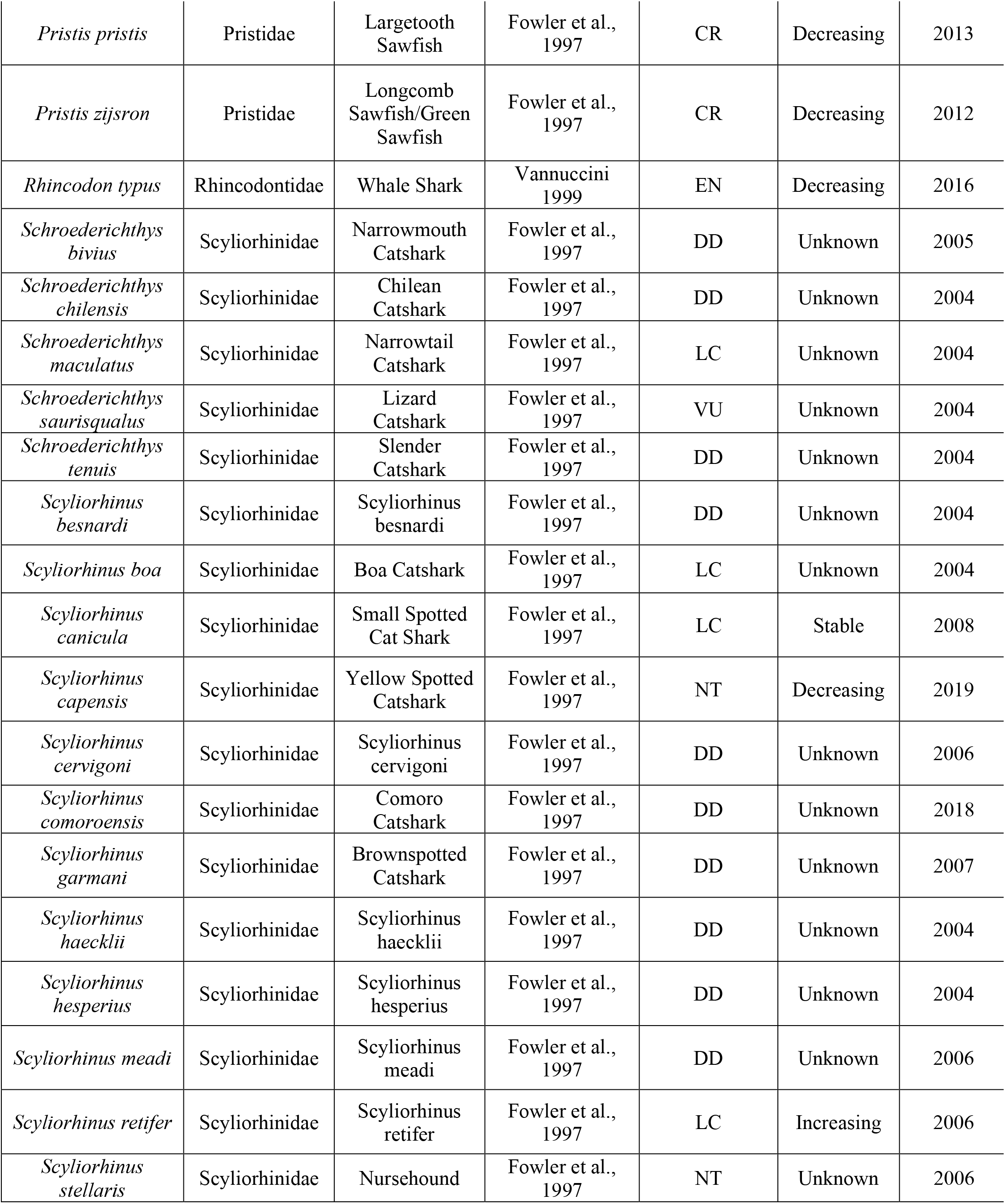
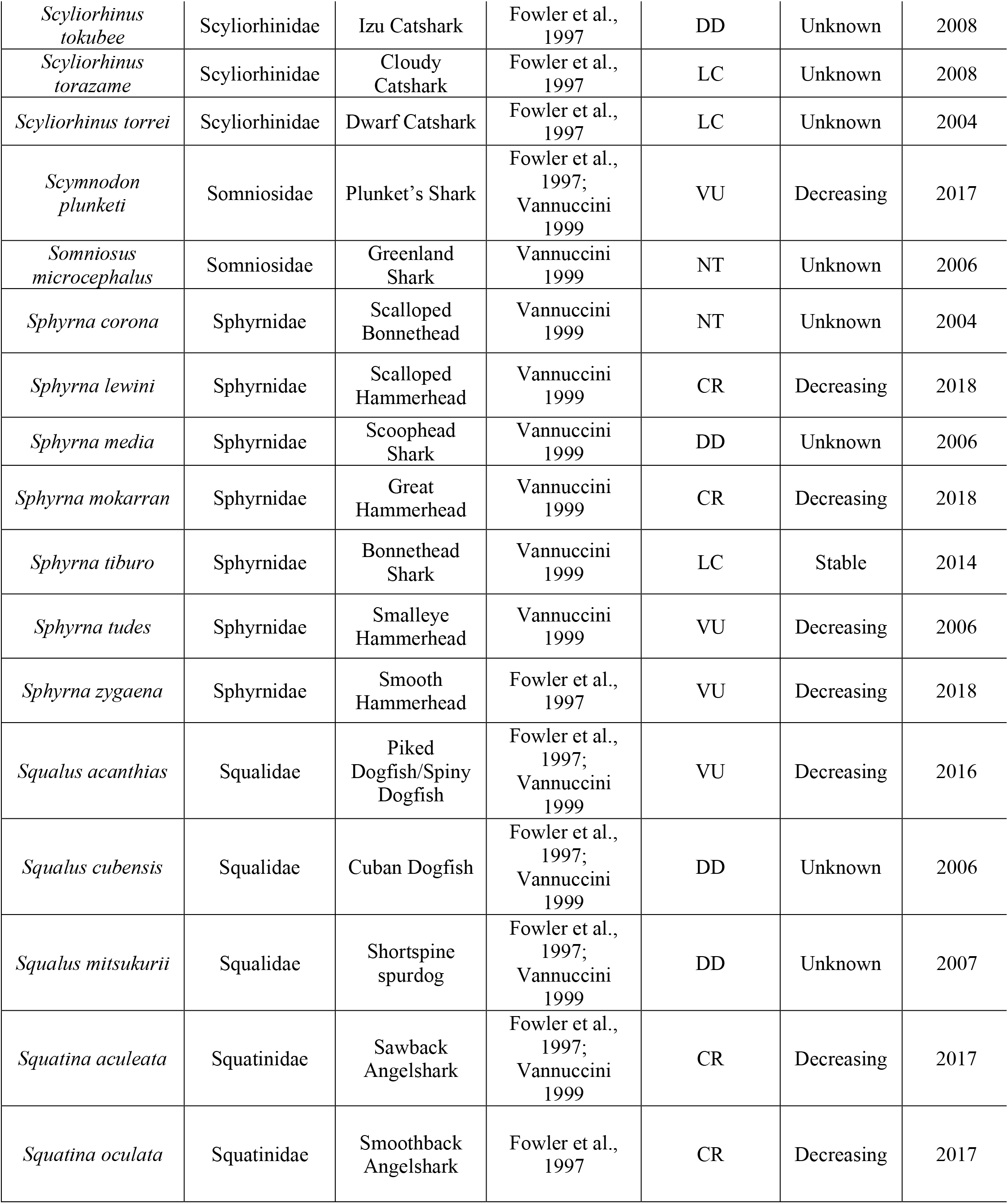
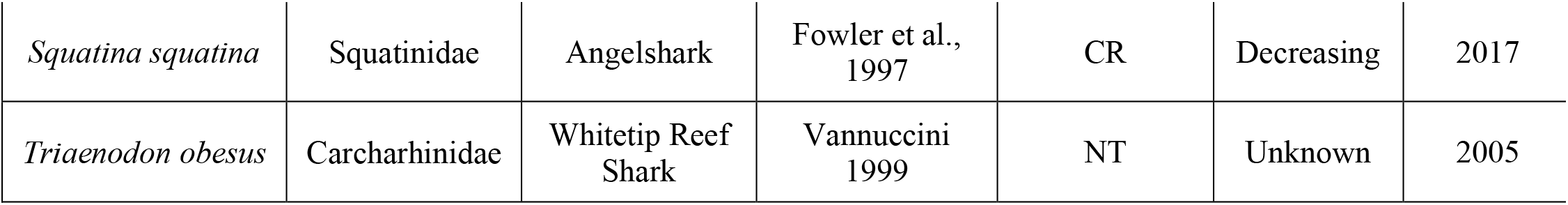

